# Discriminative Bayesian Serology: Counting Without Cutoffs

**DOI:** 10.1101/2020.07.14.202150

**Authors:** Murray Christian, Ben Murrell

**Affiliations:** Department of Microbiology, Tumor and Cell Biology, Karolinska Institutet, Stockholm 171 77, Sweden

## Abstract

During the emergence of a pandemic, we need to estimate the prevalence of a disease using serological assays whose characterization is incomplete, relying on limited validation data. This introduces uncertainty for which we need to account.

In our treatment, the data take the form of continuous assay measurements of antibody response to antigens (eg. ELISA), and fall into two groups. The *training data* includes the confirmed positive or negative infection status for each sample. The *population data* includes only the assay measurements, and is assumed to be a random sample from the population from which we estimate the seroprevalence.

We use the training data to model the relationship between assay values and infection status, capturing both individual-level uncertainty in infection status, as well as uncertainty due to limited training data. We then estimate the posterior distribution over population prevalence, additionally capturing uncertainty due to finite samples.

Finally, we introduce a means to pool information over successive time points, using a Gaussian process, which dramatically reduces the variance of our estimates.

The methodological approach we here describe was developed to support the longitudinal characterization of the seroprevalence of COVID-19 in Stockholm, Sweden.

## 1 Introduction

One approach to estimate prevalence from assay measurements begins by setting a cut-off value; samples whose measurement exceeds the value are diagnosed as infected, those below as uninfected. Rather than setting a cut-off, we model the probability that a sample is infected, given their measurement, and use this to estimate prevalence.

To illustrate our approach, we begin with the following problem. Suppose we have training data, a prior probability that a randomly drawn individual from the population is infected, and an assay measurement for such an individual. Given this information, how can we produce a posterior probability that they are infected?

A simple approach would be to fit a logistic regression [1] to the training data, and use this to calculate the probability of infection conditioned on the individual’s measurement. This approach has two drawbacks. Firstly, it neglects the prior probability that an individual is infected. Secondly, logistic regression is sensitive to the ratio of positive to negative samples in the training data, and in general this may be very different from the prevalence in the population. Our approach overcomes these issues by modelling the likelihood ratio of a measurement given the two possible infection statuses. Using Bayes’ theorem, this leads naturally to a *class-balance adjustment* in the logistic regression, which accounts for mismatch in the prevalence of infected cases in the training data and the population.

### 1.1 Assumptions

The principal assumption of our model is that the log-likelihood ratio of a measurement given possible infection status is a linear function of the measurements. Suppose that the assay measures the antibody response to *k* antigens, and let *y*_∗_ = (*y*_∗1_, …, *y*_∗*k*_) be the measurements of a random individual. Let *κ*_∗_ be their unknown infection status, coded as one for positive, zero for negative. Our model stipulates that

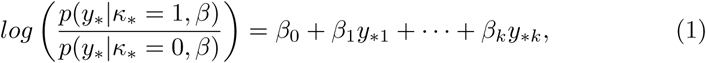

where *β* = (*β*_0_, …, *β*_*k*_) are coefficients to be estimated. Here and in what follows we abuse notation by writing

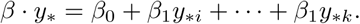

A key property of (1) is that it determines *p*(*y*_∗_ |*κ*_∗_ = 1, *β*) and *p*(*y*_∗_ |*κ*_∗_ = 0, *β*) up to a common constant that, for all downstream analyses, can be ignored. Whenever we are required to evaluate a likelihood containing these terms, we will be able to use the next two equations, which follow from (1), instead:

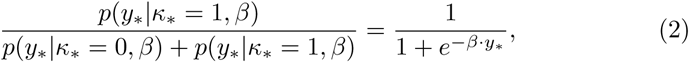

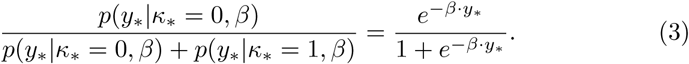

For fixed *β*, the denominator on the left side of these equations depends only on the data, and can therefore be discarded in an unnormalised posterior distribution.

### 1.2 Learning *β*s

Bayes’ theorem [2] suggests a method to estimate the coefficients *β* with a class-balance adjusted logistic regression. In terms of prior and posterior odds, Bayes’ theorem reads

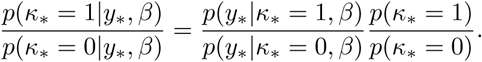

Taking logarithms, rearranging, and substituting (1) for the log likelihood-ratio, we get

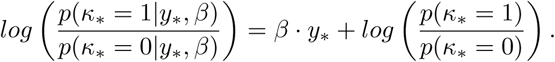

This resembles a logistic regression, with the addition of the rightmost term, the unknown log prior odds. Considered in the context of a training data set, in which the infection status of each sample is known, a natural interpretation of the prior odds is the ratio of positive cases to negative cases. Making this assumption defines a likelihood from which we can estimate the coefficients *β*.

To formally describe the logistic regression, we begin with some notation. Let *x* = (*x*_*ij*_) be the *m × k* matrix of training data, where *x*_*ij*_ is the assay measurement of sample *i* against antigen *j*. We use a single subscript to denote all the measurements for one sample, so *x*_*i*_ is the *i*^*th*^ row of *x*. Let 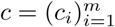 be the vector of confirmed infection statuses. Motivated by the preceding discussion, our regression model is

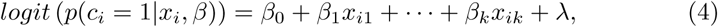

where the class-balance adjustment is the log of the ratio of frequencies in the training data:

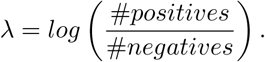

The likelihood of the training data under this model is

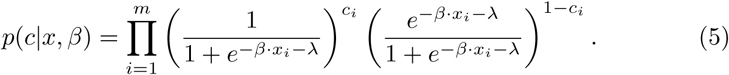

We use Bayesian inference to estimate posterior distributions over the *β* coefficients, using MCMC [3]. The uncertainty in the relationship between assay measurements and infection status is captured by the variance of these distributions.

### 1.3 Estimating the posterior probability of a single infection

Returning to our problem, we consider a randomly sampled individual with assay measurements *y*_∗_ and unknown infection status *κ*_∗_. We want an estimate for the probability that *κ*_∗_ = 1, given *y*_∗_ and the training data (*x, c*), which incorporates the prior probability of infection *p*(*κ*_∗_ = 1).

We start with

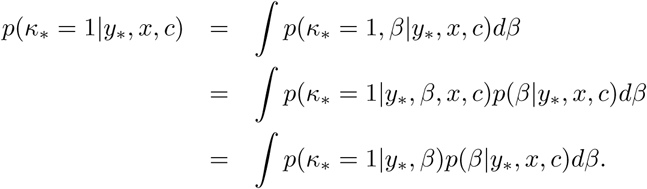

We additionally assume that all information about *β* must come from the training data, and that we can learn nothing more from unlabelled data^1^:

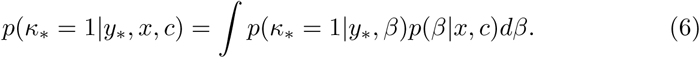

We estimate the integral (6) with an average

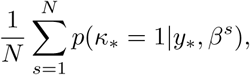

where *N* is large. The samples 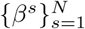 are drawn from the posterior distribution over *β*, given the training data, with the model described in section 1.2. The probabilities *p*(*κ*_∗_ = 1|*y*_∗_, *β*^*s*^) are calculated using Bayes’ theorem to write

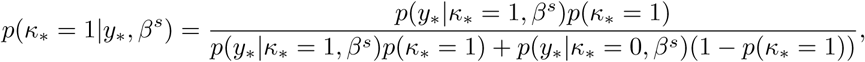

and then using (2), (3) to substitute for *p*(*y*_∗_ | *κ*_∗_ = 1, *β*^*s*^) and *p*(*y*_∗_ | *κ*_∗_ = 0, *β*^*s*^). Note how the unknown denominators in (2) and (3) fall out of the calculation.

## 2 Estimating prevalence at one point in time

Throughout this section we assume that a set of posterior beta samples has been obtained, but we omit the dependence on *β* in the notation.

Let *y* = (*y*_*ij*_) be the *n × k* population data matrix, where row *y*_*i*_ consists of the antibody responses of sample *i* to each of *k* proteins. Let *κ*_*i*_ be the unknown infection status of sample *i*, and set *κ* = (*κ*_1_, …, *κ*_*n*_). We wish to estimate *θ*, the proportion of the population that is infected.

We assume that an individual’s measurement *y*_*i*_ is independent of *θ*, given their infection status *κ*_*i*_. The joint posterior distribution for *κ* and *θ* is

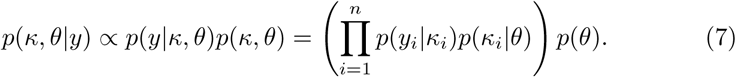

Our prior over *θ* and *κ* is

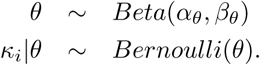

We do not model the distributions of *y*_*i*_ | *κ*_*i*_, but rather the likelihood ratio over possible infection status:

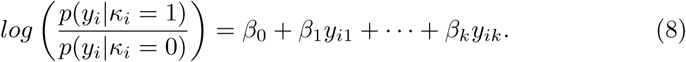

As discussed in section 1.1, this determines *p*(*y*_*i*_|*κ*_*i*_ = 1) and *p*(*y*_*i*_|*κ*_*i*_ = 0) up to a common constant that depends only on the data *y*_*i*_ (and fixed *β*). Therefore, we can evaluate the unnormalised posterior distribution on the right side of (7), and sample from the posterior distribution using MCMC techniques.

### Implementation

We estimate *θ*, marginalising out the latent variables *κ*.^2^ For concreteness, we are estimating the posterior distribution

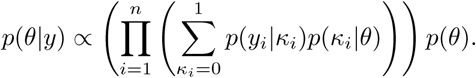

We:

1. Draw samples *β*^1^, …, *β*^*N*^ from the posterior distribution over *β* given the training data, using the likelihood (5) and weakly informative, normal priors.
2. For each posterior sample *β*^*s*^, draw one sample *θ*^*s*^ from the posterior distribution *p*(*θ* | *y*) (which depends on *β*^*s*^). To do this we run *K* iterations of Metropolis’ algorithm to sample from *p*(*θ* | *y*), and set *θ*^*s*^ to the last sample obtained.

The set 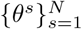 represents a sample from the posterior distribution for *θ*, from which moments and Bayesian intervals [4] can be computed.

## 3 Estimating prevalence at multiple times

We extend the model to estimate the prevalence of the disease from samples that have been collected at multiple times. Our method assumes that seroprevalence is monotonically increasing, which requires that antibodies in infected individuals wane at a rate slower than the accumulation of new infections^3^. If antibody waning is negligible over the time scales considered, then we can interpret sero-prevalence as the total proportion of individuals that are currently infected or that have been infected in the past.

Suppose that we have samples from times *t*_1_, …, *t*_*w*_. Let

- *y*_*t*_ = (*y*_*t,ij*_) be the *n*_*t*_ × *k* matrix of assay measurements for the *j*^*th*^ protein of the *i*^*th*^ donor at time *t*, whose *i*^*th*^ row we write as *y*_*t,i*_,
- let *κ*_*t*_ = (*κ*_*t*,1_, …, *κ*_*t,n*_*t*) be latent variables representing the infection status of donor *i* at time *t*, and
- let *θ*_1_, …, *θ*_*w*_ be the prevalence at times *t*_1_, …, *t*_*w*_.

We assume that the (*y*_*t*_, *κ*_*t*_) are independent of all other variables in the model, given *θ*_*t*_. For each time *t*, the model for the variables (*y*_*t*_, *κ*_*t*_) | *θ*_*t*_ is the same as described in the single time case:

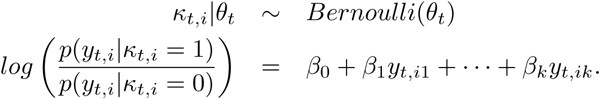

We introduce a prior over the initial prevalence, and over the increase in prevalence over time. Each increase is the product of an unknown exposure probability *η*_*i*_ and the proportion of the population that has not yet been infected. We use a Gaussian process prior over the logit of the exposures, allowing flexibility in the prevalence trajectories. Formally, we let

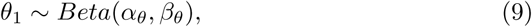

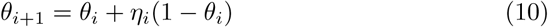

for *i* = 1, …, *w* − 1, and

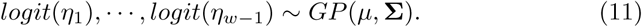

Prevalence increases monotonically and is bounded above by one.

The Gaussian process (11) is realized at constant time intervals (eg. weeks). The covariance terms in **Σ** are determined by a rational quadratic kernel, which is a mixture of squared exponential kernels of varying time scales

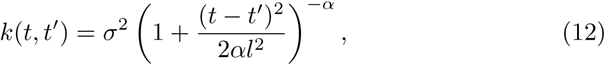

*σ*^2^ is the variance, *l*^−2^ is the average time scale, and *α* is a parameter that controls the distribution of time scales [5]. See Figure 1 for draws from our

**Figure 1:**
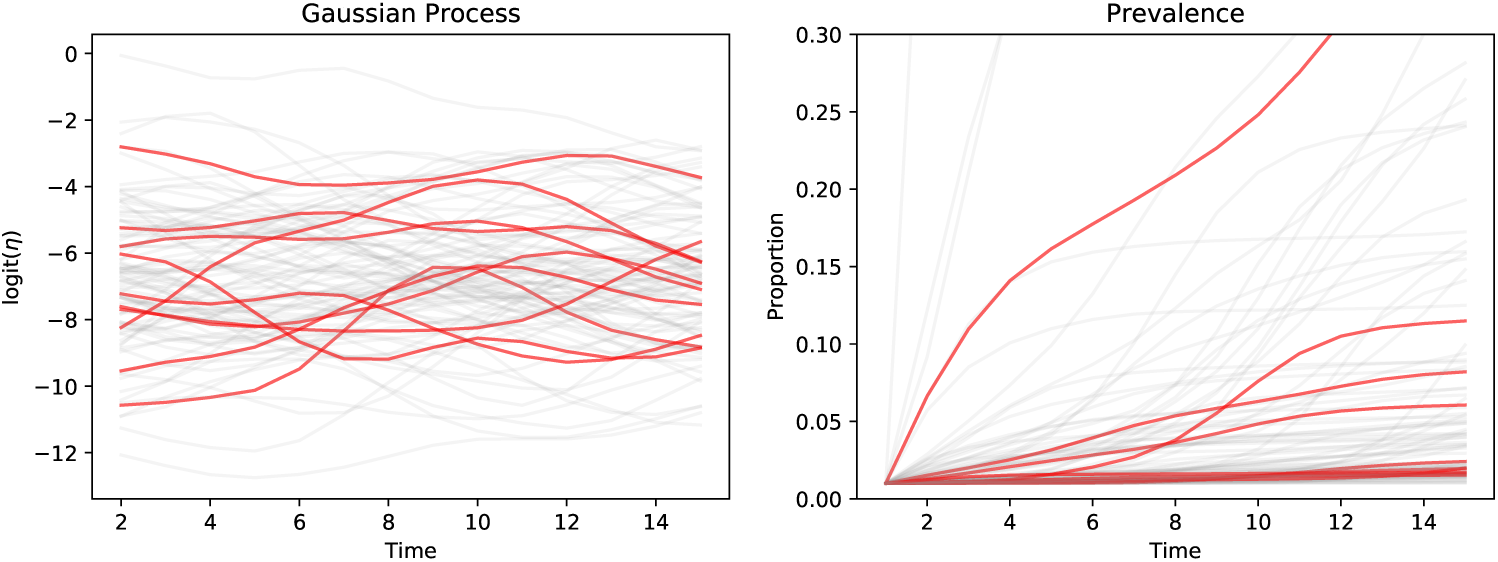
Curves drawn from the Gaussian process prior (left) and corresponding prevalence curves (right), starting from the same initial prevalence. One hundred curves are drawn in grey to illustrate the distribution of curves under the model. In red, ten trajectories have been highlighted. The process parameter values are described in the text.

### Implementation

We estimate the posterior distribution over all the model parameters except the latent states *κ*_*t,i*_, which we marginalise out. The posterior distribution is

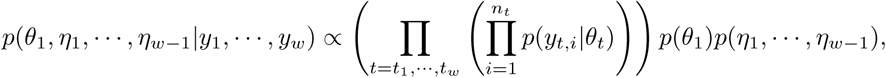

Where

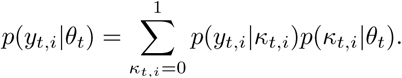

Our implementation is now an extension of the one described in the single time case:

1. Draw samples *β*^1^, …, *β*^*N*^ from the posterior distribution over *β* given the training data, using the likelihood (5), and weakly informative, normal priors over the *β* parameters.
2. For each posterior sample *β*^*s*^, draw one sample 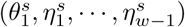, from the posterior distribution over these parameters (which depends on *β*^s^). To do this we run many iterations of Metropolis’ algorithm to sample from *p*(*θ*_1_, *η*_1_, …, *η*_*w*−1_|*y*_1_, …, *y*_*w*_), and keep the last sample obtained. (Proposals to the parameters (*η*_1_, …, *η*_*w*−1_) are made by adding a draw from a mean-zero Gaussian process to the current values.)

Our sample from the posterior distribution over prevalences (*θ*_1_, …, *θ*_*w*_) is 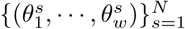. We refer to each of these samples as a posterior prevalence trajectory.

## 4 Simulations

In our simulations we consider a single ELISA measurement (*k* = 1), drawn from either of two Gamma distributions, which correspond to positive and negative cases. The components of the simulations which we vary are, (*i*) the degree of overlap between positive and negative distributions, (*ii*) the number of training samples and the class-balance, and (*iii*) the number of population samples.

Firstly, we examine three degrees of overlap between the positive and negative distributions, using a Gamma(2, 0.1) negative distribution throughout, and Gamma(5, 0.1), Gamma(10, 0.1) and Gamma(30, 0.1) positive distributions^4^ for the *large overlap, slight overlap* and *well-separated* settings, respectively (see Figure 2).

**Figure 2:**
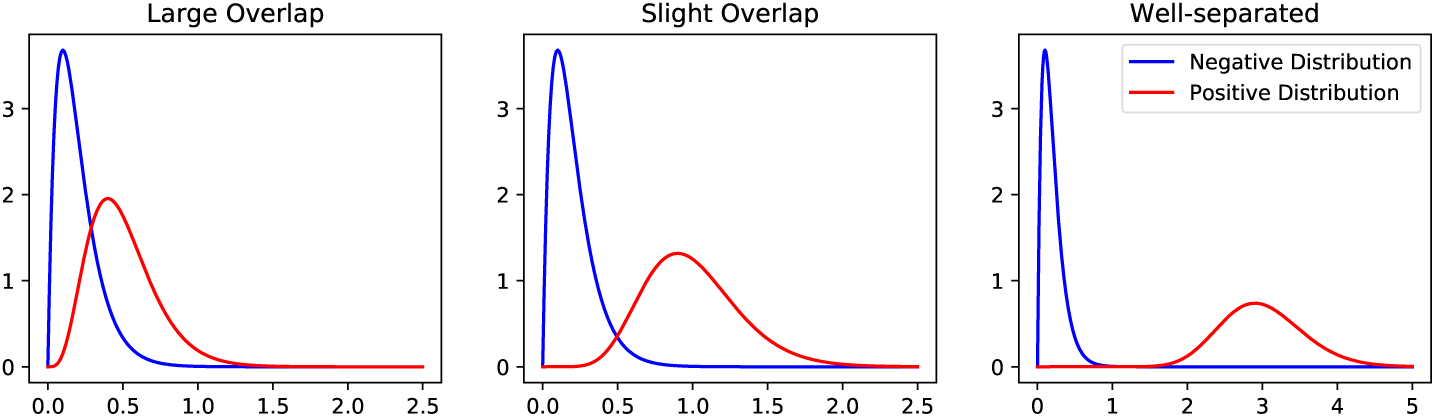
Three degrees of overlap between positive and negative Gamma distributions.

Secondly, the number of training samples in the positive and negative classes are divided into five settings, two of which skew the class-balance by a factor of 20:

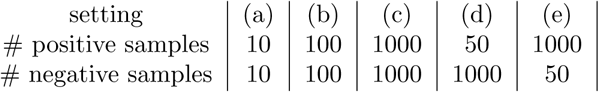

Thirdly, we take the number of population data samples obtained at each time to be 40, 200 or 1000.

For each of the 45 settings just described, we run ten replicate simulations, and the training data, sample times, true prevalence sequences and population data are drawn new in each replicate.

### Data generation

To generate true prevalence sequences that mimic plausible real-world scenarios, we ran a Susceptible-Infected-Recovered (SIR) model in which the basic reproduction number *R*_0_ is subject to small stochastic changes. We then drew a random sample of times, and set the true prevalence at each of these times to be the proportion of the population that is either infected or recovered; see Figure 3.

**Figure 3:**
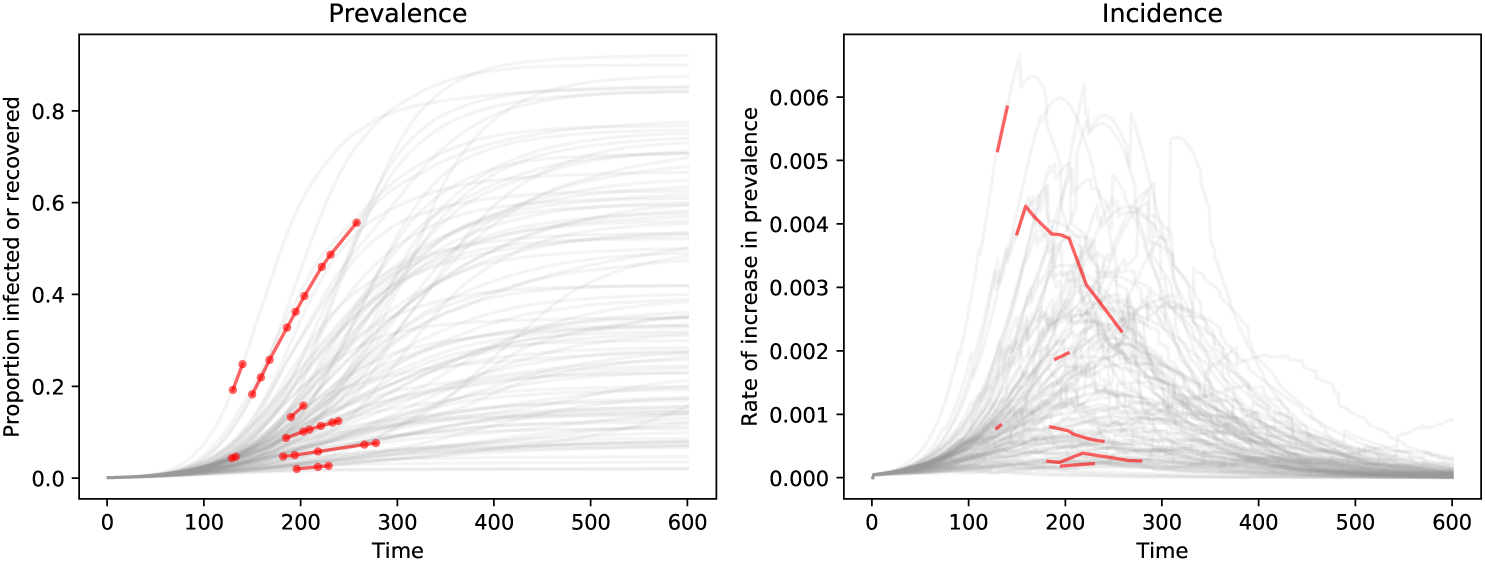
Prevalence curves (left) and corresponding incidence curves (right) drawn from our SIR model. One hundred curves are drawn in grey to illustrate the distribution of curves under the model. In red, seven trajectories have been highlighted, with randomly drawn sample times indicated on the prevalence curves.

Given the sample times, and the sequence of true prevalence values at these times, we draw binomial positive case counts from these, and then simulate ELISA measurements from the positive and negative distributions for each case in the corresponding class.

### Model settings for inference

In each replicate, we ran inference under our model with the following settings. Our priors on the beta coefficients were *β*_0_, *β*_1_ ∼ *Normal*(0, 20). For the Gaussian process prior over prevalence curves we used

- a mean vector *µ* with all entries equal to −7, and
- a covariance matrix **Σ** determined by a rational quadratic kernel with *σ*^2^ = 2, *l* = 5 and *α* = 0.5; see (12).

The mean vector *µ* controls the expected slope of the corresponding prevalence curve, and the kernel parameters control the variance in the slope (*σ*^2^) and frequency of fluctuations in the slope (*l* and *α*). We selected the parameter values by inspecting trajectories simulated under the prior, ensuring that the process represents a reasonable distribution over prevalence curves; see Figure 1.

In each replicate we ran the model with a Beta(1, 15) and a Uniform(0, 1) prior over the initial prevalence *θ*_1_.

### 4.1 Results

The results of our simulations show that the model performs well in the following key respects:

- it is sufficiently flexible to estimate non-linear trends in prevalence,
- it appropriately captures uncertainty in the posterior estimates,
- its posterior estimates are accurate,
- it is robust to a skewed class-balance in the training data, and
- it is not sensitive to choice of prior on *θ*_1_.

### Model flexibility

The prevalence during a pandemic is a non-linear function of time, and linear functions will not always yield satisfactory approximations, especially over large time scales. Indeed, under a standard SIR model, the prevalence, *I* + *R*, grows exponentially at first, and eventually plateaus as the total proportion that will become infected is saturated. The examples in Figure 4 demonstrate that our model is capable of learning non-linear prevalence functions.

**Figure 4:**
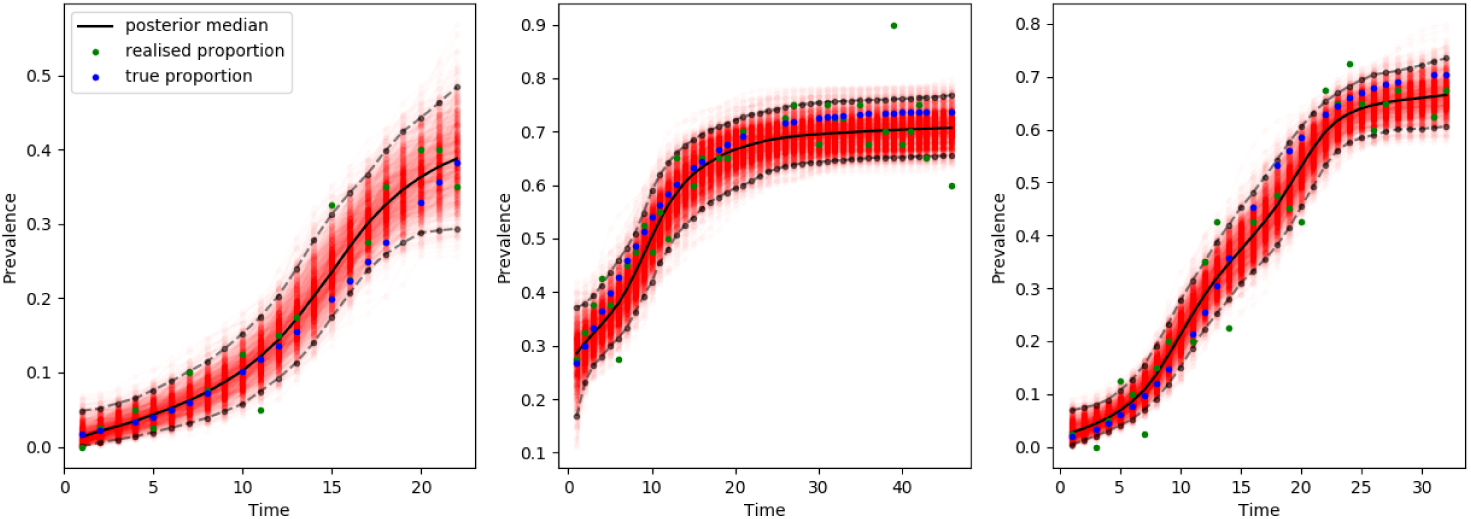
Non-linear prevalence trajectories over large time scales. Each plot shows 1000 posterior prevalence trajectories (in red), with their median (in black) and 95% credible interval (dashed). The true prevalence, and the realised proportion of positive samples at each sampled time, are also shown. In these simulations, 40 population samples were drawn at each of 18, 33 and 25 times, from slightly overlapping or well-separated positive and negative distributions.

### Uncertainty in posterior estimates

When we infer prevalence, there are several sources of uncertainty that should affect the confidence in the posterior estimates. The three sources we examine here are the degree of overlap in the positive and negative distributions, the number of training samples, and the number of population samples.

SI Figure 1 illustrates the decrease in width of the posterior intervals with increase in the separation between positive and negative distributions, and in the number of training examples, whilst holding the size of the population data roughly constant. SI Figure 2 illustrates the tendency for the width of the posterior intervals to decrease as the size of the population data increases, either by increasing the number of samples at each time, or the number of sampling times.

### Accuracy of estimates

To assess the accuracy of posterior estimates of prevalence, we defined a *simulation error* as 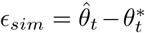, where *t* is a random sample time in that simulation, 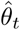 is the median posterior sample of prevalence at time *t*, and 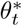 is the true prevalence. In Figure 5 we plot the distribution of simulation errors over all simulations, and a breakdown by degree of overlap in the positive and negative distributions. In the large overlap case, the errors are often undesirably large, but the accuracy improves as the degree of overlap decreases.

**Figure 5:**
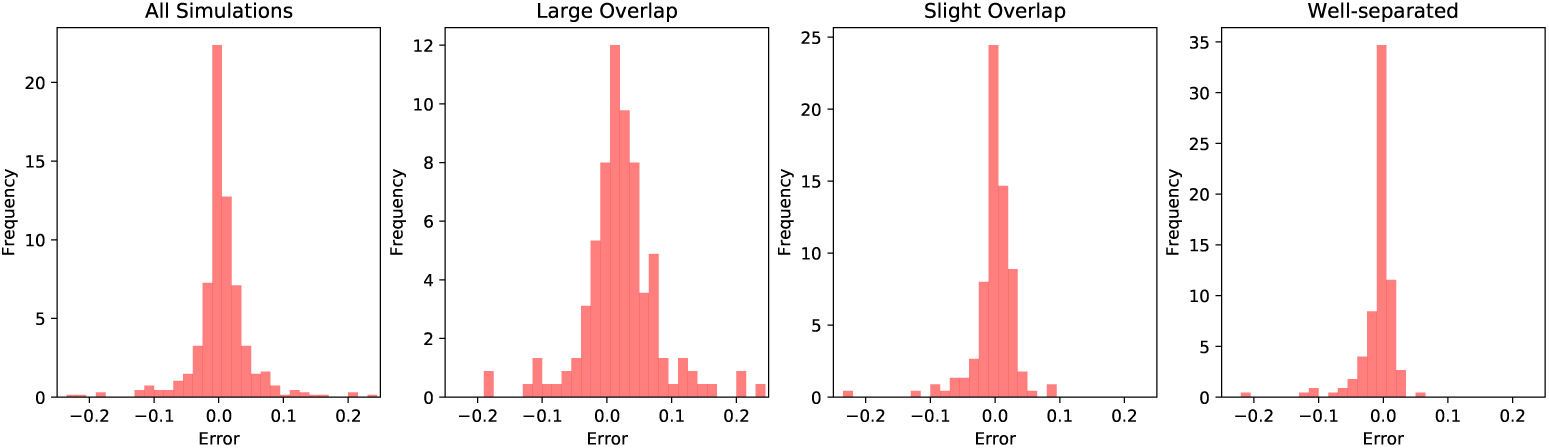
The distribution of errors *E*_*sim*_ for all simulations (left), and for all simulations with a given degree of overlap between positive and negative distributions (right).

### Robustness under skewed class-balance

The model is able to accommodate a twenty-fold imbalance between the positive and negative classes in the training data, provided that the overlap between positive and negative distributions is not large. Figure 6 compares the errors under balanced training data to each of the scenarios with unbalanced training data, demonstrating that accuracy is not affected by a skewed class-balance.

**Figure 6:**
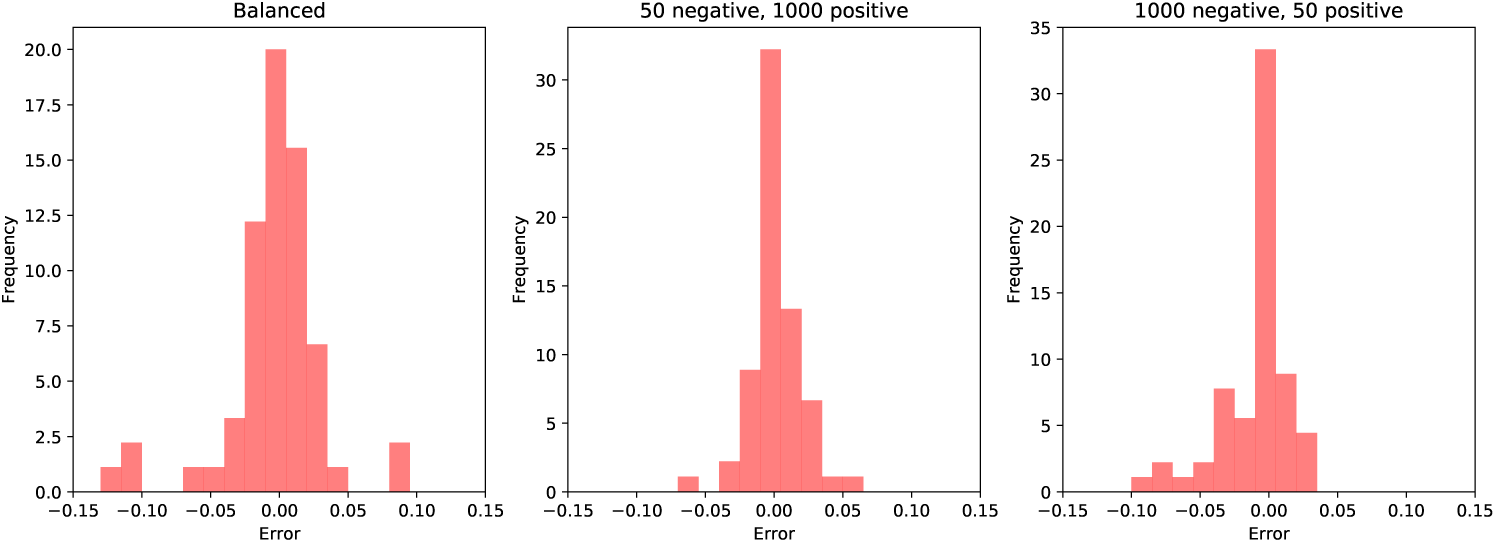
Distribution of errors under balanced (left) and imbalanced (right) training data. Simulations with a large overlap in the positive and negative distributions are not included here, and the data for the balanced case further excludes simulations with ten training samples in each class.

### Effect of the prior on *θ*_1_

To examine the effect of our choice of prior over *θ*_1_, we ran our model on the data from each simulation, using a Beta(1, 15) prior and a flat Uniform(0, 1) prior. Inspection of the posterior distributions shows almost no difference between the two cases. Furthermore, Figure 7 shows that the distribution of median posterior *θ*_1_ estimates over all simulations is the same for both choices of prior. Therefore the model’s estimates should be insensitive to any reasonable specification of prior on *θ*_1_.

**Figure 7:**
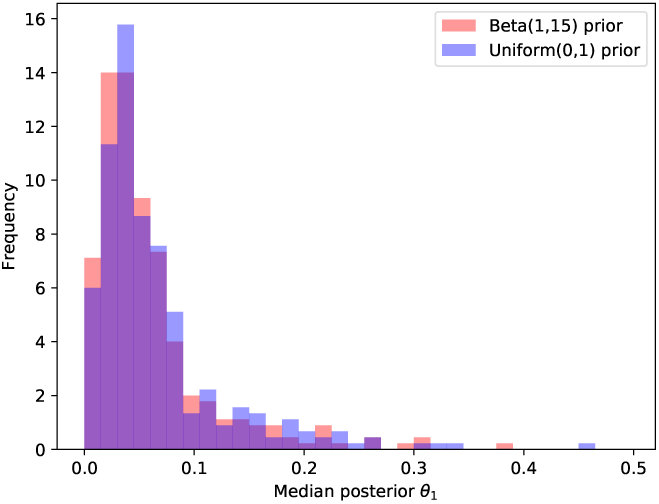
Histograms of median posterior *θ*_1_ estimates for all simulations, in the case of Beta(1, 15) and Uniform(0, 1) priors over *θ*_1_.

## 5 Discussion

We began by mentioning that standard serological diagnostics use a single threshold to decide between positive and negative cases, and then estimate how frequently this is correct in two directions: sensitivity and specificity. It is possible to apply principled adjustments to these hard-threshold models when estimating population prevalence, correcting for the expected proportion of false positives and false negatives [6]. But the underlying hard-threshold model treats a sample whose measurement is just over the threshold as having the same uncertainty as a strong positive signal. Here we have sought to introduce a principled way of relaxing such a hard threshold, while accounting for multiple relevant sources of uncertainty.

The core of the approach presented here is that we can exploit a discriminative model of infection given assay measurement to estimate population prevalence. We use logistic regression to model infection status, but logistic regression can be replaced with any discriminative probabilistic model of infection probability given assay measurement. For logistic regression, the class balance adjustment propagates into the linear term (as *λ* in eq. (4)), but this isn’t a general requirement, and this class balance adjustment can be introduced to any discriminative model with probabilistic outputs.

We additionally show that pooling information over time provides large gains in the precision of prevalence estimates when multiple time points are sampled sparsely. One avenue for extension would be to implement hyperparameter inference for the Gaussian Process, which would become progressively more useful as the number of sampling time points increases.

Our transformed Gaussian Process prior over new infections is a modular component, and more sophisticated pooling strategies that encode epidemic dynamics may be required when modeling prevalence over longer time scales. This would provide efficient pooling, while allowing us to infer epidemic parameters (eg. *R*(*t*)) from serology data, but would have to be sufficiently flexible to avoid becoming over-constrained by model assumptions.

While we do not demonstrate this, covariates of infection probability can be included in the model, and their coefficients estimated.

Finally, future work will require comparison to emerging approaches that either correct for uncertain sensitivity and specificity when estimating sero-prevalence [6], or that use cutoff-free approaches, but in a generative modeling framework [7].

## Acknowledgements

We thank Xaquín Castro Dopico and Gunilla Karlsson Hedestam for contributing to the framing of this problem originally, and we thank Chris Wallace and Nastasiya Grinberg for many helpful comments and discussions.

---

**SI Figure 1:**
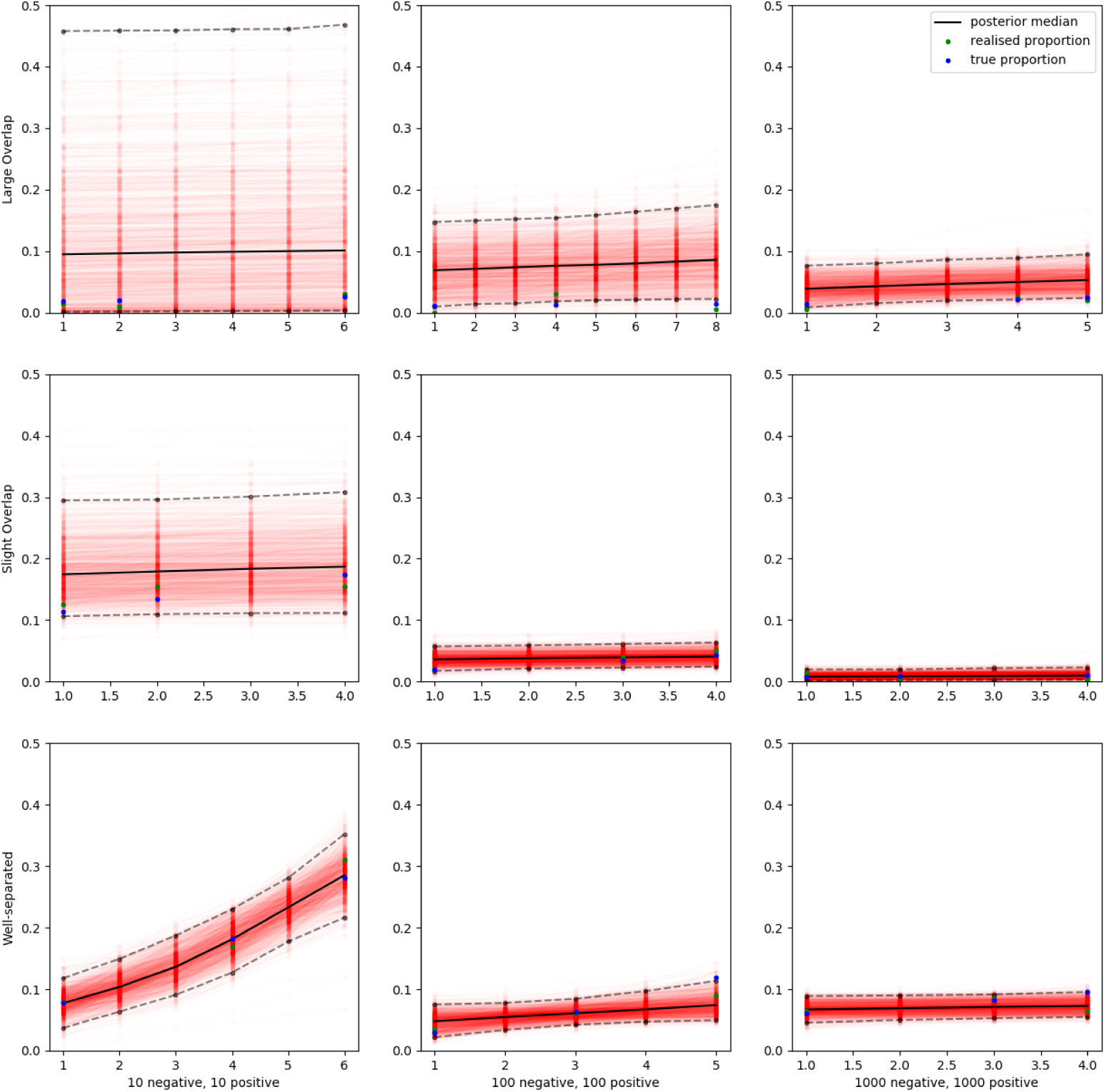
Posterior prevalence trajectories under three degrees of overlap of the positive and negative distributions (top to bottom row), and three sizes of the training samples (left to right column). Each simulation pictured here had population data comprised of 200 samples from three or four times.

**SI Figure 2:**
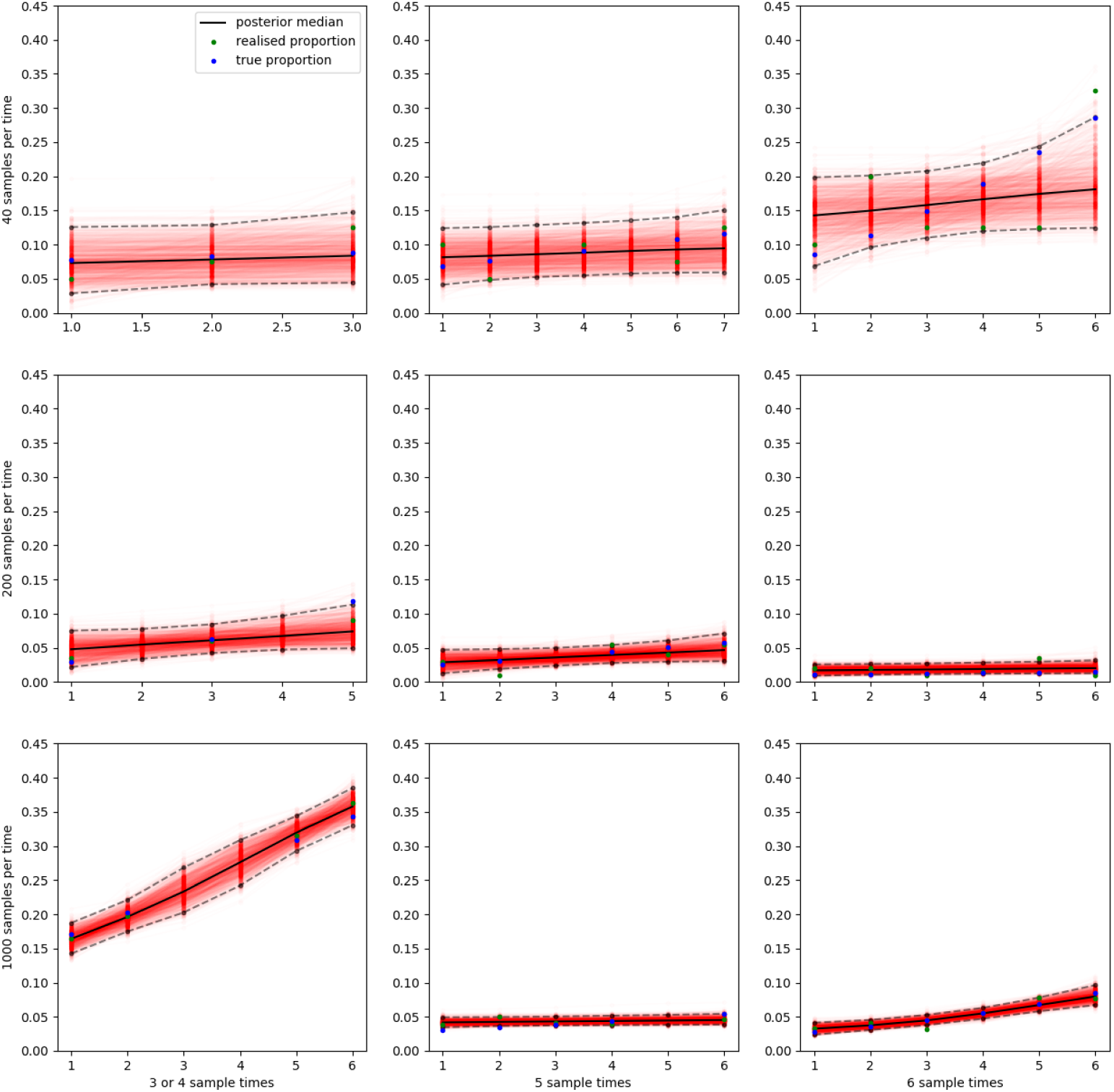
Posterior prevalence trajectories with increasing population data, due to more samples at each time (top to bottom), and sampling at more times (left to right). Each simulation pictured here was drawn with low overlap between positive and negative distributions, and training data comprised of 100 positive and 100 negative samples.

## Footnotes

1 By not explicitly modeling the distribution of positive and negative classes we are forced to give up the possibility of refining our estimates of the infection probabilities from any unlabelled data. This would be possible under a full generative model, but likely overly sensitive to assumptions about the distribution of positive and negative cases.

2 If one is interested in each individual’s posterior probability of infection, one can jointly sample *κ* and *θ*.

3 This monotonicity can be relaxed at the expense of increased variance in the seroprevalence estimates

4 We parametrise Gamma distributions by *shape* and *scale*, whose product gives the mean.

## References

[1] Cox D.R. (1958). The regression analysis of binary sequences. J. Royal Statistical Society. Series B (Methodological) Vol. 20, No. 2, pp. 215–242.

[2] Bayes T. (1763). An essay toward solving a problem in the doctrine of chances Phil. Trans. Roy. Soc. 330–418.

[3] Metropolis N., Rosenbluth A.W., Rosenbluth M.N. and Teller A.H. (1953). Equation of state calculations by fast computing machines. J. Chem. Phys. 21, 1087.

[4] Jaynes E.T. and Kempthorne O. (1976) Confidence Intervals vs Bayesian Intervals. In: Harper W.L., Hooker C.A. (eds) Foundations of Probability Theory, Statistical Inference, and Statistical Theories of Science. Springer.

[5] Rasmussen C.E. and Williams C.K.I. (2006). Gaussian Processes for Machine Learning. MIT Press.

[6] Gelman A. and Carpenter B. (2020). Bayesian analysis of tests with unknown specificity and sensitivity. https://doi.org/10.1101/2020.05.22.20108944

[7] Bouman J.A., Bonhoeffer S. and Regoes R. (2020) Estimating seroprevalence with imperfect serological tests: exploiting cutoff-free approaches. https://doi.org/10.1101/2020.04.29.068999

